# Stress induces divergent gene expression among lateral habenula efferent pathways

**DOI:** 10.1101/2020.05.15.098319

**Authors:** Marjorie R. Levinstein, Kevin R. Coffey, Russell G. Marx, Atom J. Lesiak, John F. Neumaier

**Affiliations:** Graduate Program in Neuroscience, University of Washington, Seattle, WA, 98104, USA; Department of Psychiatry and Behavioral Sciences, University of Washington, Seattle, WA, 98104, USA; Department of Genome Sciences, University of Washington, Seattle, WA, 98104, USA; Department of Pharmacology, University of Washington, Seattle, WA, 98104, USA

## Abstract

The lateral habenula (LHb) integrates critical information regarding aversive stimuli that shapes decision making and behavioral responses. The three major LHb outputs innervate dorsal raphe nucleus (DRN), ventral tegmental area (VTA), and the rostromedial tegmental nucleus (RMTg). LHb neurons that project to these targets are segregated and nonoverlapping, and this led us to consider whether they have distinct molecular phenotypes and adaptations to stress exposure. In order to capture a time-locked profile of gene expression after repeated forced swim stress, we used intersectional expression of RiboTag in rat LHb neurons and next-gen RNA sequencing to interrogate the RNAs actively undergoing translation from each of these pathways. The “translatome” in the neurons comprising these pathways was similar at baseline, but diverged after stress, especially in the neurons projecting to the RMTg. Using weighted gene co-expression network analysis, we found one module comprising genes downregulated after stress in the RMTg-projecting LHb neurons; there was an overrepresentation of genes associated with phosphoinositide 3 kinase (PI3K) signaling in this module. Reduced PI3K signaling in RMTg-projecting LHb neurons may be a compensatory adaptation that alters the functional balance of LHb outputs to GABAergic vs. monoaminergic neurons following repeated stress exposure.

---

Stress disorders, anxiety, and depression are associated with altered functional connectivity of key brain regions and are critical determinants of associated symptoms ^1^, including the lateral habenula (LHb), a small nucleus in the epithalamus that acts as an anti-reward nucleus. While it receives inputs from diverse regions throughout the brain, it has three main efferent pathways – to the ventral tegmental area (VTA), the rostromedial tegmental nucleus (RMTg), and the dorsal raphe nucleus (DRN) ^2,3^. These three outputs contribute to different elements of adaptive responses to stress ^3,4^. The neurons comprising these three main output pathways appear to be quite segregated as they do not send collaterals to more than one of these targets ^5-8^. LHb neurons are mainly glutamatergic, contributing excitatory synapses onto both GABAergic and serotonergic neurons in the DRN at roughly equivalent rates and directly onto dopaminergic neurons in the VTA ^9,10^. LHb afferents to RMTg are also excitatory, and in turn RMTg inhibits both the DRN and VTA through GABAergic projection neurons. Both the DRN and VTA send reciprocal projections to each other and back to the LHb, and serotonin and dopamine released from these projections modulate LHb excitability ^11-15^. Thus, there is a complex interplay of excitation and inhibition of DRN and VTA that is controlled by the relative activation of these LHb output pathways.

The LHb may be an important target for treating stress disorders ^2,16,17^. Stress exposure activates LHb neurons intensely and induces c-Fos, a marker of neuronal activity ^18^. Stimulation of LHb neurons promotes passive avoidance ^19,20^ while inhibition of the LHb, and its connection to both serotonergic and dopaminergic regions, reduces anxiety- and depression-like behaviors ^21-23^. Inhibition of the LHb via the inhibitory DREADD receptor hM_4_Di decreases passive coping in the forced swim test (FST), a measure of behavioral despair ^24^. LHb projections to the DRN are thought to mediate this effect ^4^.

Recently, a surge in transcriptomic data emphasizes the heterogeneity of LHb neurons. Wagner, et al. ^25^ used *in situ* hybridization images from the Allen Brain Atlas to map transcript expression throughout the LHb and its subregions, finding ten distinct anatomical patterns of expression. Recent single cell RNAseq profiles of the habenular nuclei identified several types of distinct neurons ^26,27,^ although none of these appeared to be pathway-specific. However, these studies did not explicitly examine gene expression between the three major LHb outputs. Given the heterogeneity of the LHb transcriptome and the functional differences between its anatomically segregated output pathways, it seems plausible that gene expression in these pathways respond differentially to stress. Additionally, while both Hashikawa, et al. ^27^ and Wallace, et al. ^26^ studied the murine habenula, this is the first study examining the genetic profile of LHb neurons in the rat.

To investigate this, we used next-gen sequencing after intersectional expression of RiboTag in neurons from each of these pathways to isolate ribosome-associated RNAs actively undergoing translation^28^, a method that reveals evolving changes in gene expression in response to stress ^29^. This is the first such application of intersectional translatome analysis in rat neurons and by capturing ribosome-associated RNA from the soma and dendrites, may be sensitive to rapid changes in translation following stress. We found that LHb neurons projecting to RMTg are differentially regulated by stress in comparison to those projecting to DRN or VTA.

## Results

### Read Depth

Three weeks after viral vector injections, we compared handled but unstressed rats to rats subjected to a two-day forced swim stress and then all animals were sacrificed 3 hrs after completion of the stress procedure. This differs from the conventional Porsolt Forced Swim Test, where the behavior is measured during the second swim session. RNA was purified, sequencing libraries were prepared, and samples were sequenced as described in the Materials and Methods section (Figure 1). The IP samples were sequenced to a depth of 7.4 ± 0.4 x10^6^ double stranded reads whereas the Input RNA samples were sequenced to a read depth of 1.1 ± 0.1 x10^7^ double stranded reads (Table 1). We sequenced Input RNA from pooled samples including each animal from that treatment group rather than each individual; this allowed us to estimate and subtract the small amount of contaminating input RNA that is carried forward during RiboTag immunoprecipitation ^30^.

**Table 1.** Read Depth.

| Sample Type | Pathway | Stress | n | Read Depth (mean ± SEM) |
| --- | --- | --- | --- | --- |
| IP | DRN | Stressed | 3 | $6.624 \times 10^6$ ($\pm 1.742 \times 10^6$) |
| IP | DRN | Unstressed | 3 | $8.601 \times 10^6$ ($\pm 0.958 \times 10^6$) |
| IP | RMTg | Stressed | 3 | $7.957 \times 10^6$ ($\pm 0.077 \times 10^6$) |
| IP | RMTg | Unstressed | 3 | $7.001 \times 10^6$ ($\pm 0.079 \times 10^5$) |
| IP | VTA | Stressed | 3 | $7.055 \times 10^6$ ($\pm 0.077 \times 10^6$) |
| IP | VTA | Unstressed | 3 | $7.217 \times 10^6$ ($\pm 0.202 \times 10^6$) |
| Input | DRN, RMTg, VTA | Stressed | 3 (pooled each pathway) | $11.84 \times 10^6$ ($\pm 1.667 \times 10^6$) |
| Input | DRN, RMTg, VTA | Unstressed | 3 (pooled each pathway) | $10.01 \times 10^7$ ($\pm 1.371 \times 10^6$) |
| IP | Sham | Unstressed | 3 | $5.299 \times 10^6$ ($\pm 0.773 \times 10^6$) |
| Input | Sham | Unstressed | 3 | $8.05 \times 10^6$ ($\pm 0.813 \times 10^6$) |

**Figure 1.**
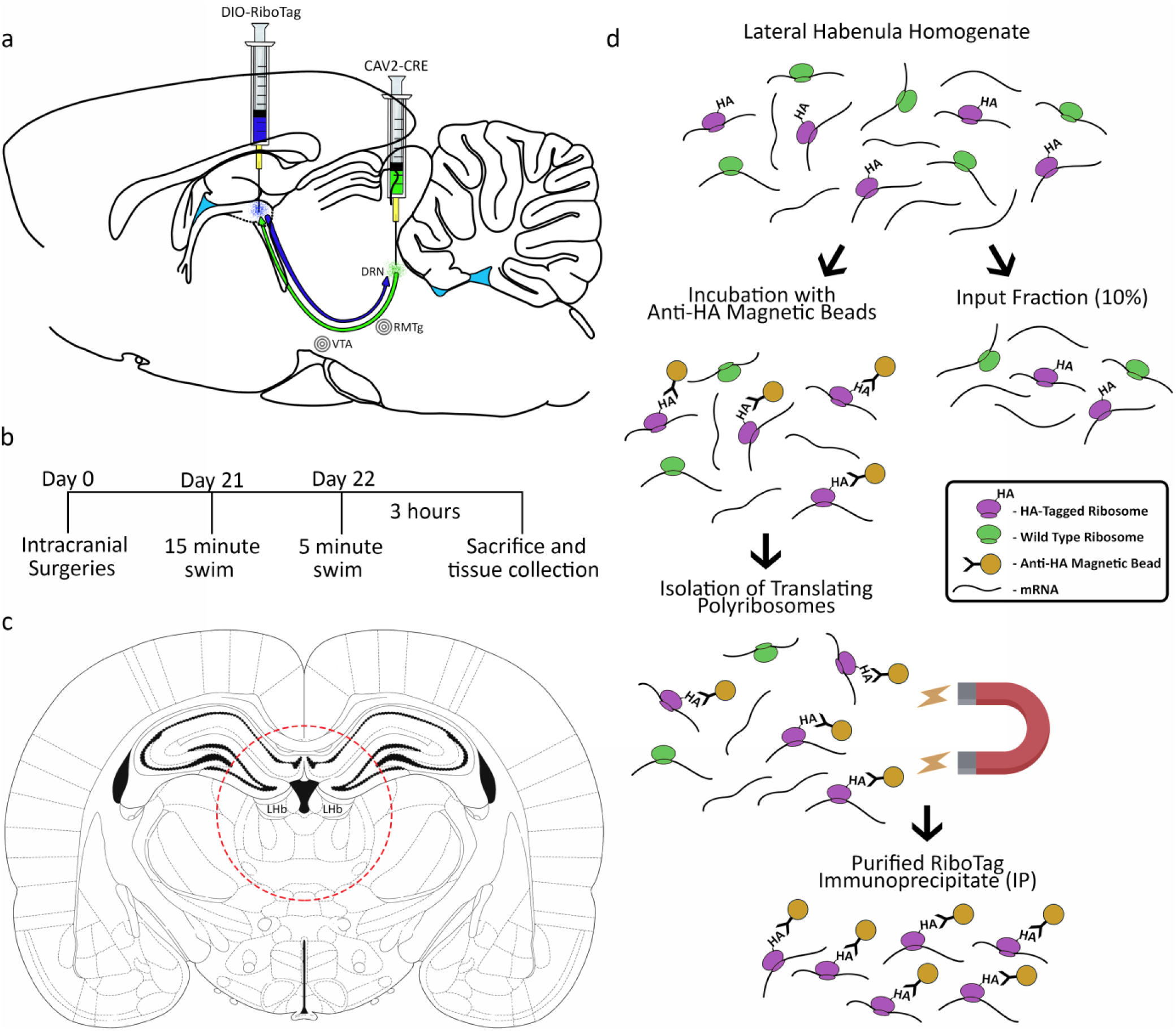
Intersectional viral mediated gene transfer and RiboTag strategy. **a)** Rats were injected with AAV8-DIO-RiboTag into the LHb and CAV2-Cre injected into one of the three target regions, either the DRN, RMTg or VTA. **b)** Experimental timeline. **c)** Location of tissue punch for LHb extraction. **d)** RiboTag immunoprecipitation protocol.

### Stress Induced Divergent Gene Expression in LHb Output Pathways

Initially, we compared differential gene expression in the three pathways using pairwise comparisons among the unstressed conditions (usDRN, usVTA, usRMTg), then the stressed conditions (sDRN, sVTA, sRMTg), and then between stressed and unstressed conditions within each pathway using DESeq2 (Figure 2). In all cases, if the gene was more highly expressed in the first group, the Wald score is positive (upregulated) and if it was more highly expressed in the second group, the Wald score is negative (downregulated). We used an FDR of q= 0.1 for these initial pair-wise comparisons between pathways. Gene expression profiles in the LHb projection neurons were relatively similar among the three pathways under the basal, unstressed condition (Figure 2a-c). Thirty-one genes were differentially expressed between the unstressed DRN pathway and unstressed RMTg pathway (Figure 2a). Of these, 21 genes were more highly expressed in the DRN pathway. Eleven genes were differentially expressed between DRN and VTA-projecting LHb neurons (Figure 2b). The VTA and RMTg pathways showed the greatest difference at baseline, with 236 DEGs (Figure 2c). Most of these genes (141) were expressed at greater levels in the VTA pathway compared to the RMTg pathway.

**Figure 2.**
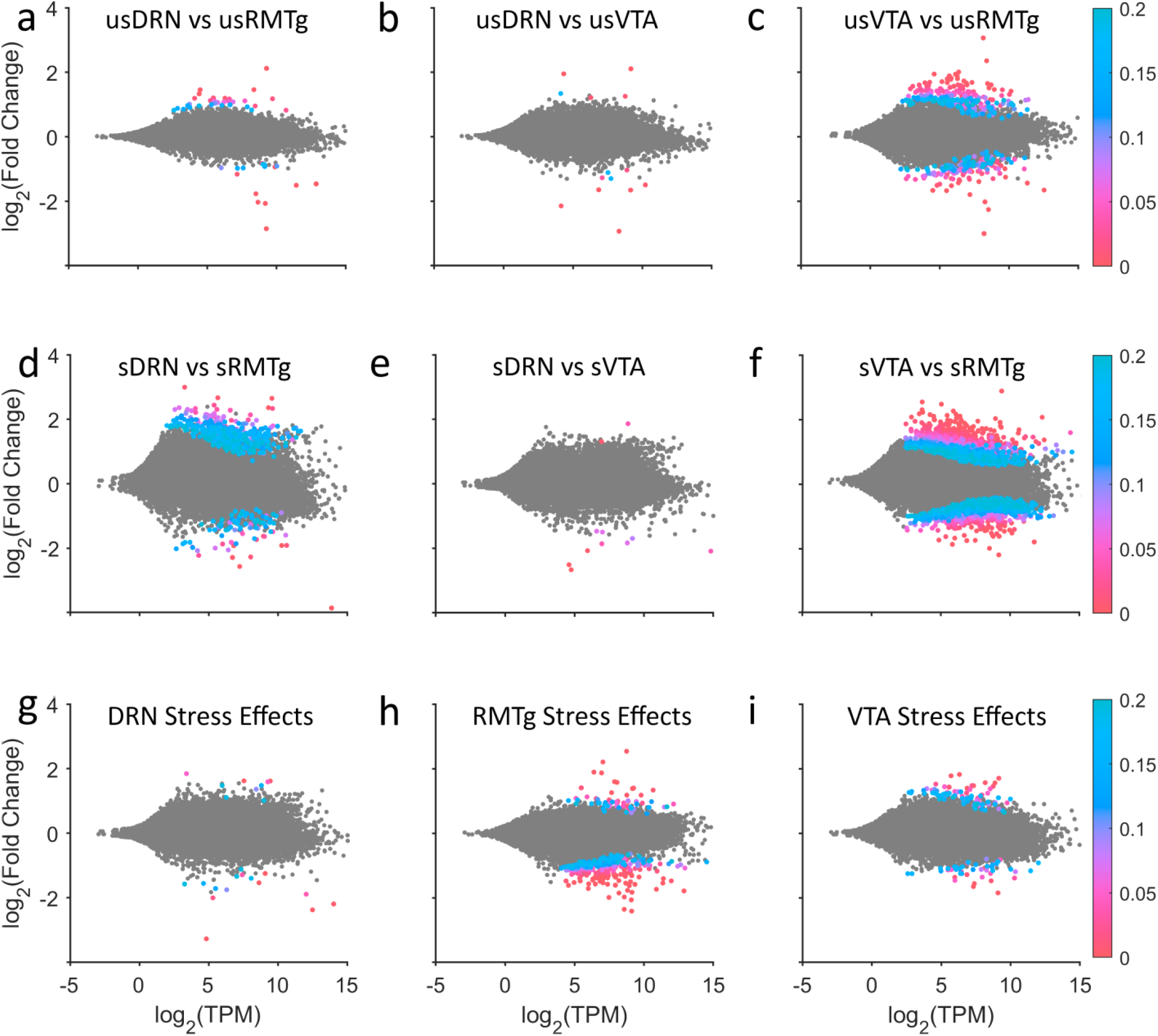
Pairwise DESeq2 comparisons. The top row shows comparisons between the unstressed pathways **(a-c)**; there were few differences between LHb neurons projecting to the DRN and neurons projecting to the RMTg or VTA, although several hundred genes differed between the RMTg and VTA pathways at baseline. The middle row shows comparisons between the stressed pathways **(d-f)**. There were only a few differences between the DRN and the VTA pathway, but stress exposure caused dramatic gene expression changes in the RMTg-projecting neurons relative to either the DRN or VTA pathways. The bottom row shows stress effects within each pathway **(g-i)**. As is evident, the neurons projecting to the RMTg had the most differentially expressed genes, and most of those were downregulated.

Interestingly, gene expression in stressed rats showed striking differences between the pathways (Figure 2d-f). 125 genes were differentially expressed between the stressed DRN and RMTg pathways, and 95 of those genes were more highly expressed in the DRN pathway (Figure 2d). As with the unstressed comparisons, the stressed DRN and VTA pathways remained similar – only 11 genes were differentially expressed (Figure 2e). Further, 1,078 genes were differentially expressed between the stressed VTA and RMTg pathways (Figure 2f). The majority of these genes (657) were expressed at greater levels in the VTA pathway.

We next determined the effects of stress within the neurons of each pathway (Figure 2g-i). Stress had the smallest impact on gene expression within the DRN-projecting LHb neurons–only 14 genes were different with 5 being upregulated and 9 being downregulated following stress (Figure 2g). In the RMTg-projecting neurons, 204 genes were changed by stress; with 38 being upregulated and 166 being downregulated (Figure 2h). 63 genes were changed by stress in the VTA pathway – 40 were upregulated and 23 were downregulated (Figures 2i).

Figure 3 illustrates overlapping DEGs between and within the pathways. When comparing the effects of stress on each pathway, there were only a small number of gene expression differences that overlapped across pathways. Sema4d was consistently downregulated with stress in all three pathways (Wald scores =-5.2 to −8, q score <0.001). Several other genes (Plekha7, Zfp467, and Slfn5) were significantly differentially expressed in some but not all cases (q<0.001-0.09) (Figure 3a). When comparing the unstressed pathways, only a few genes overlapped. Compared to usDRN, no genes were more heavily expressed in both the usRMTg and usVTA (Figure 3b). When compared to the usRMTg pathway, 6 were more highly expressed in both the usDRN and usVTA (Figure 3c). When comparing to the usVTA pathway, Trpc4 was more highly expressed in both usDRN and usRMTg pathways (usDRN vs usVTA Wald= 4.7, q=0.006; usVTA vs usRMTg Wald=-3.4, q=0.06) (Figure 3d). Oxytocin was positively differentially expressed in the DRN in both of these comparisons (usDRN vs usRMTg Wald=4.93, q=0.0011; usDRN vs usVTA Wald=5.63, q<0.0001) (Figure 3c-d). Compared to the sDRN pathway, Hcfc1 (sDRN vs sRMTg Wald=-8.4, q<0.0001; sDRN vs sVTA Wald=-4.4, q=0.04) and AABR07037536.1 (sDRN vs sRMTg Wald=-3.4, q=0.09; sDRN vs sVTA Wald=-5.7, q=0.0002) were more heavily expressed in the sRMTg and sVTA pathways (Figure 3e). While there were only a handful of overlapping DEGs between the pairwise comparisons within stressed and unstressed cases, 52 genes were more heavily expressed in both the stressed DRN and VTA pathways than in the stressed RMTg pathway (Figure 3f). When compared to the sVTA pathway, no genes were more heavily expressed in both the sDRN and sRMTg pathways (Figure 3g).

**Figure 3.**
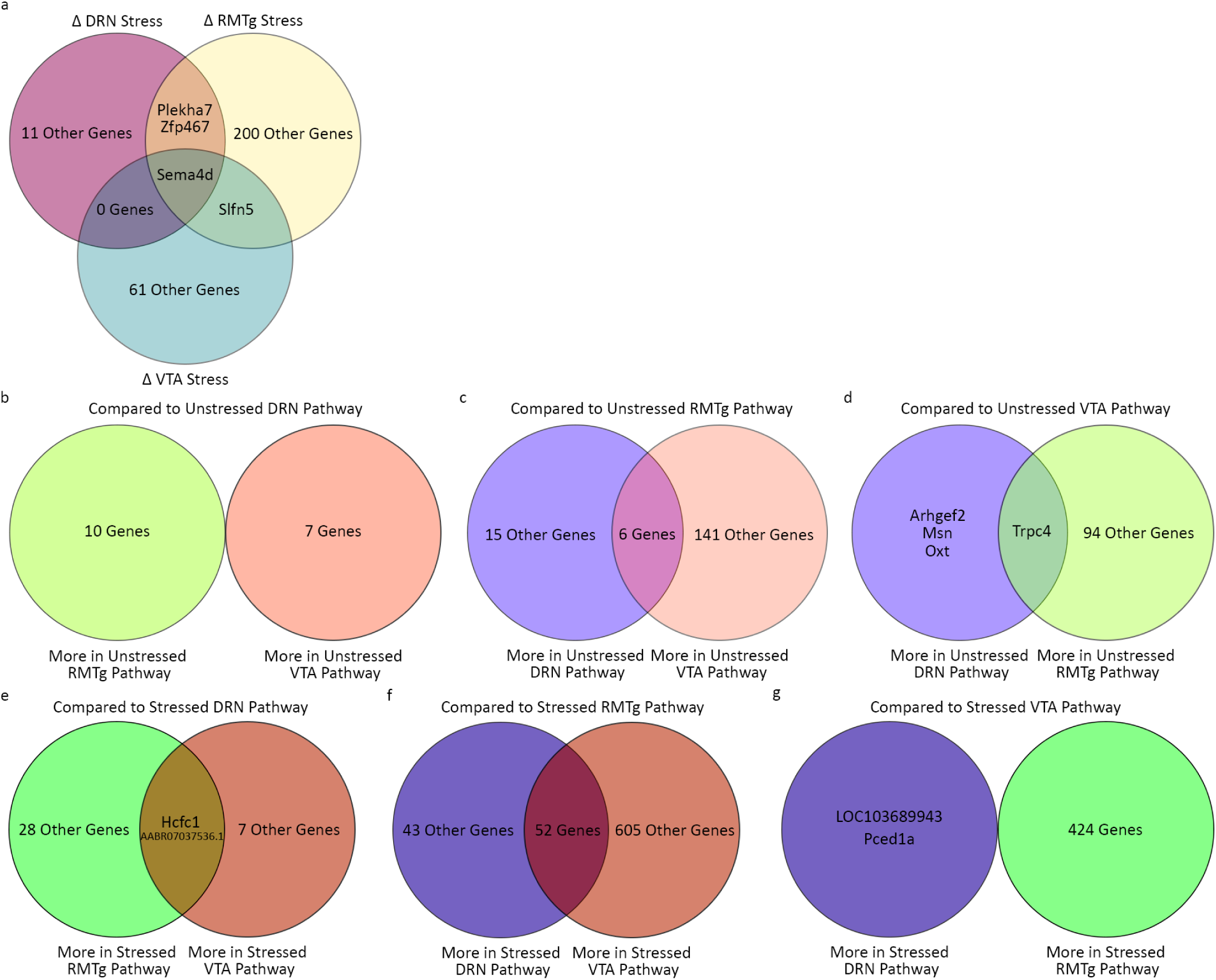
Venn diagrams of differentially expressed genes (q<0.1). **a)** The intersection of DEGs in stressed vs. unstressed neurons in each pathway were few; Sema4d was the only gene to be downregulated after stress in all pathways. **b-d)** Pairwise comparisons between RNA from the unstressed conditions revealed few differences between the pathways although the DRN and VTA pathways differed the least from each other and shared the most DEGs relative to RMTg. **e-g)** RNA in the RMTg-projecting neurons from stressed animals diverged from the other pathways. While the RNA from the DRN and VTA pathways of stressed animals were similar to one another, RNA in the RMTg-projecting neurons had ∼4.5 fold more DEGs than in the comparisons from unstressed animals.

### Stress Produced Opposite Patterns of Gene Set Enrichment in DRN and VTA Pathways as Compared to the RMTg Pathway

Next, we used a curated gene set strategy to evaluate whether there were patterns of changes in sets of related genes using WebGestalt to perform Gene Set Enrichment Analysis (GSEA). The significantly enriched gene sets were similar across different databases and showed a clear pattern: in most cases stress led to a reduction in expression of a particular gene set within the DRN (sDRN vs usDRN) and VTA (sVTA vs usVTA) pathways, but an increase within RMTg (sRMTg vs usRMTg); in the few cases where this pattern was not replicated, stress induced changes in RMTg were always in the opposite direction than in VTA or DRN (Figure 4). While the function-defined gene sets shared this consistent pattern, gene sets relating to microRNA or transcription factor targets were not prominent nor shared across pathways and are not presented.

**Figure 4.**
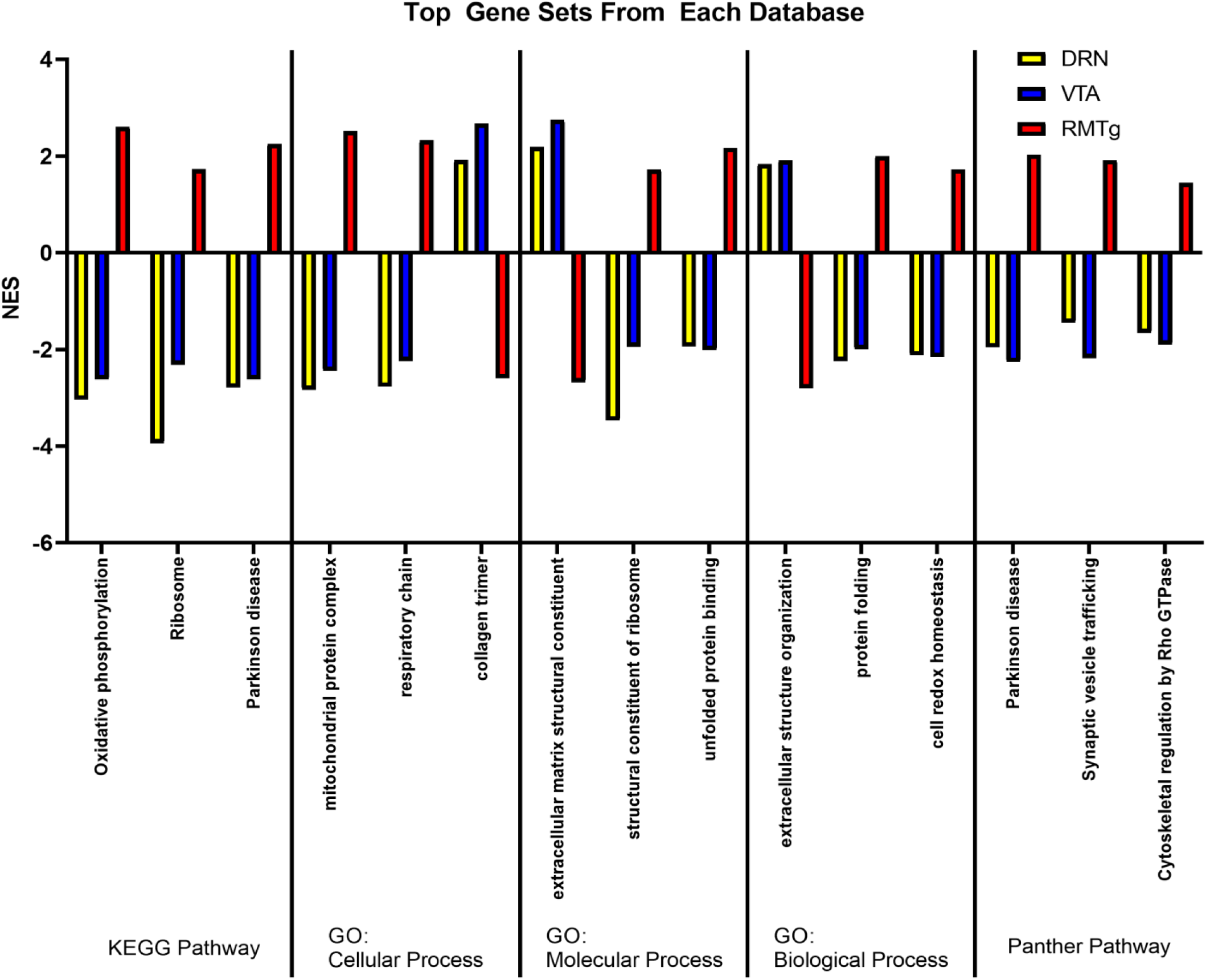
Gene set enrichment analysis. The average normalized enrichment scores (NES) for stressed vs. unstressed animals for each pathway of the three gene sets found in all three pairwise comparisons with the largest differences are illustrated for each data base queried. Of note, in each gene set, the direction of RMTg changes (i.e. upregulation or downregulation of DEGs) was in the opposite direction as for the DRN and VTA pathways. Unlike other analyses in this paper, GSEA mostly detected gene sets that were upregulated by stress in the RMTg pathway. *GO - Gene Ontology*

### Stress Downregulated a PI3-Kinase Related Gene Network in the RMTg Pathway

Lastly, we performed Weighted Gene Co-expression Network Analysis (WGCNA), an unguided topographical method for identifying modules of genes with correlated expression. Initially, WGCNA identified 96 modules; however, many of the module eigengenes were highly correlated. Modules were merged down to 35 using hierarchical clustering with a tree cut height of 1.5. The module colors (Figure 5a) and q-scores for every gene for each of the nine pairwise comparisons are mapped onto minimum spanning trees in Figure 5. Notably, while the unstressed pathway comparisons (Figure 5b-d), and even the within-pathway stress comparisons (Figure 5h-j) have relatively few low q-values, the stressed DRN vs stressed RMTg (Figure 5e) and stressed RMTg vs stressed VTA (Figure 5g) contained many genes with low q-values as illustrated in the pseudocolor scale. Many of these DEGs map to the “Gold” module. Additionally, the distribution of Wald scores for each pair-wise comparison in each module are presented as violin plots in Figure 6. The Gold module stood out with an average Wald score >±2 in multiple comparisons. The Gold module reached this threshold in the Stressed vs. Unstressed RMTg pathway (average Wald = −2.192, Figure 6b) and the Stressed RMTg vs. Stressed VTA pathway (average Wald = −2.317, not shown). Additionally, this module reached a less stringent threshold of >±1.5 in the Stressed RMTg vs. Stressed DRN pathway comparison (average Wald = −1.584, not shown).

**Figure 5.**
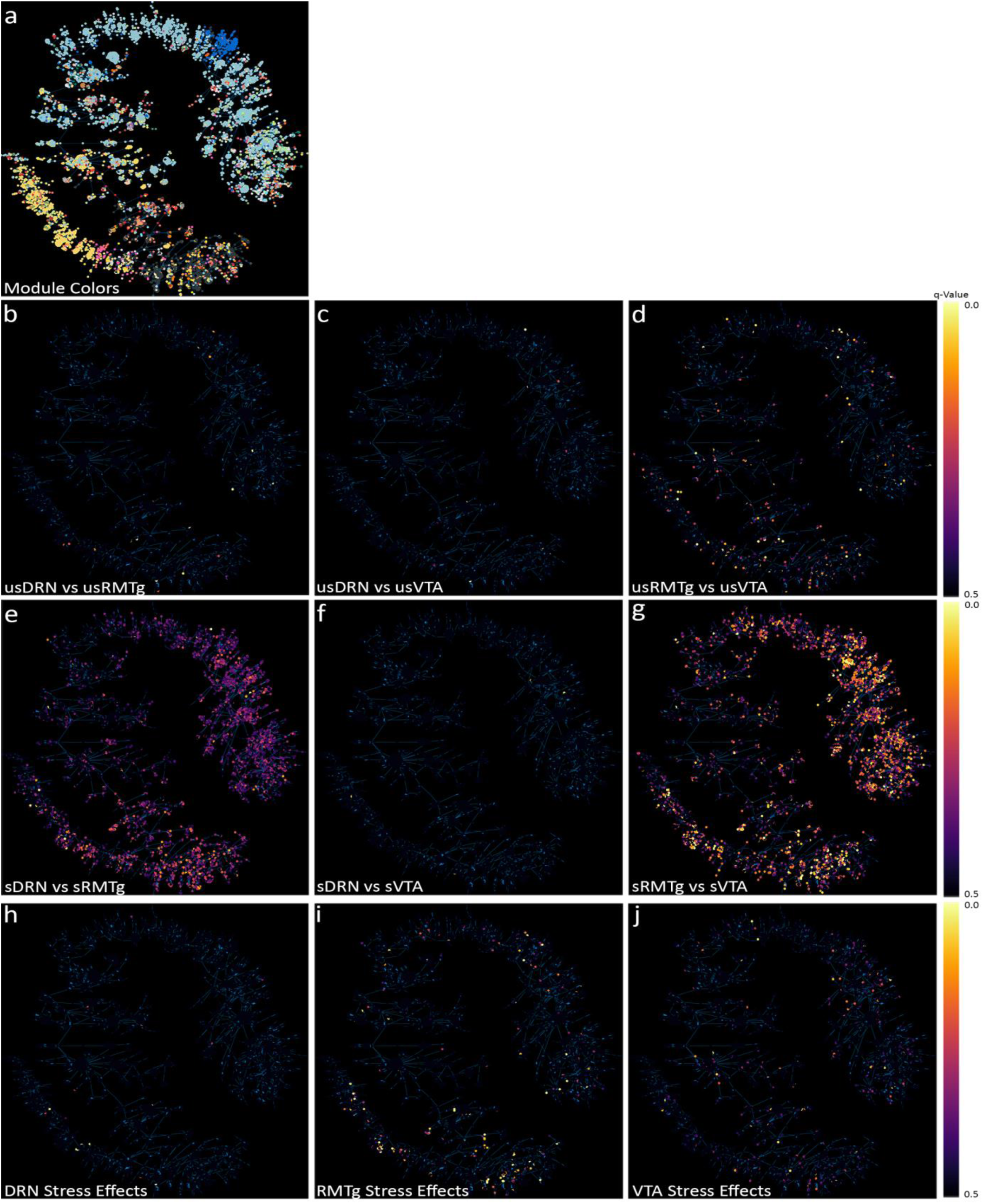
Minimum spanning trees of WGCNA results. **a**) Two-dimensional minimum spanning trees illustrate the 35 gene modules identified by weighted gene coexpression network analysis (WGCNA). Using the original DESeq2 analysis, the q-values for each individual gene is shown for pairwise comparisons of unstressed DRN to unstressed RMTg (**b**), unstressed DRN to unstressed VTA (**c**), and unstressed RMTg to unstressed VTA (**d**). There were relatively few changes apparent between the pathways in unstressed animals, but after stress, there were larger differences apparent in in the stressed DRN vs. stressed RMTg (**e**), stressed DRN vs. stressed VTA (**f**), and stressed RMTg vs. stressed VTA (**g**). The RMTg vs. VTA showed the most significant differences. (**h-j**) shows the pairwise comparisons for stressed vs. unstressed DRN (**h**), RMTg (**i**), and VTA (**j**).

**Figure 6.**
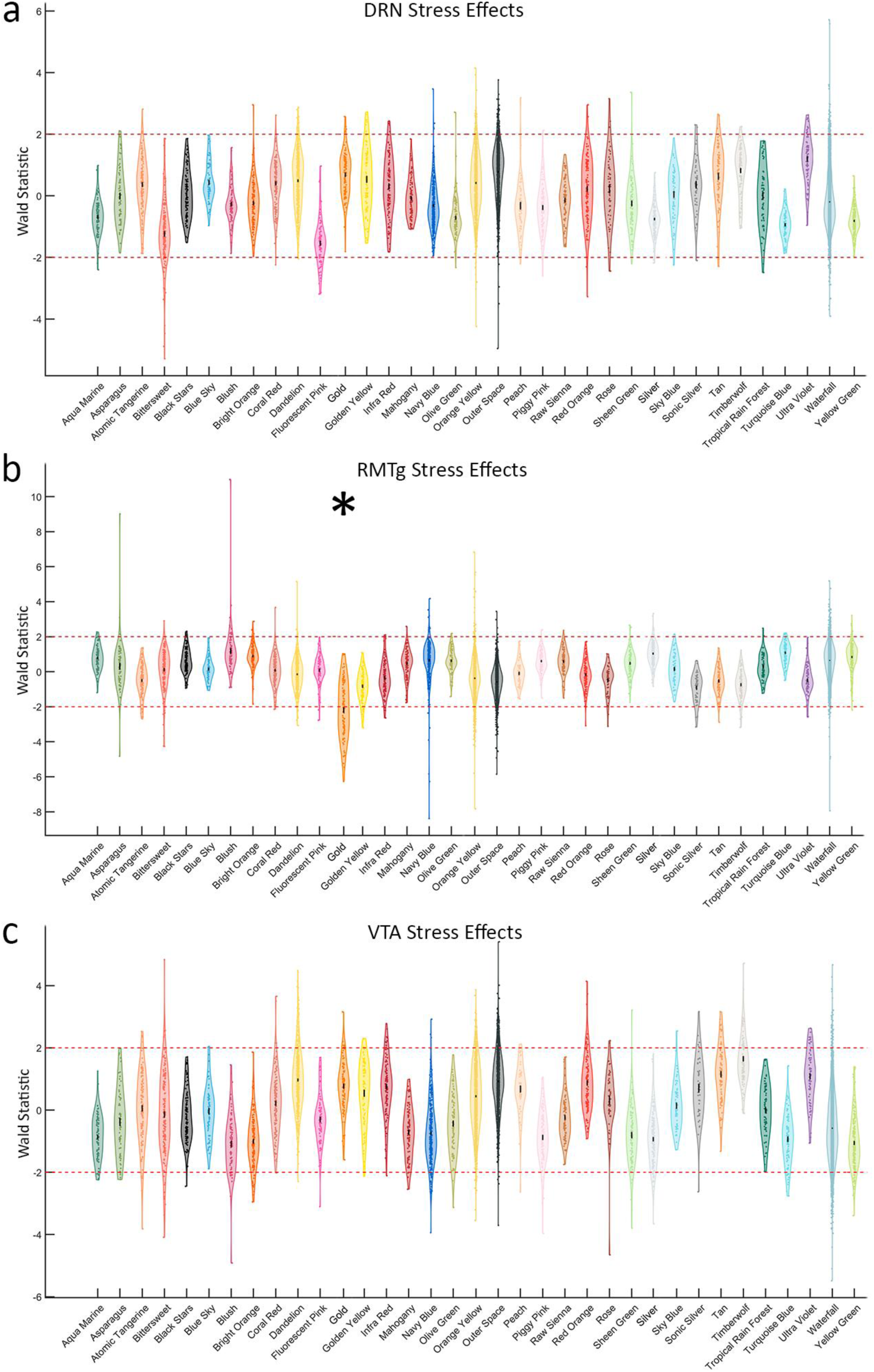
The Wald score for each gene within the 35 modules assigned by WGCNA. **a)** Stress effects in the DRN pathway. **b)** Stress effects in the RMTg pathway. The Gold module is downregulated with stress in the RMTg pathway (mean Wald= −2.192) **c)** Stress effect in the VTA pathway.

Figure 7 depicts each gene in this module’s importance within the network using a centrality metric (the more central genes have many strong bright lines); the genes most central to this module topologically are analogous to “hub” genes in other schemes. The centrality of individual genes in the Gold module correlated significantly with the Wald score in pairwise comparisons of differential expression for stressed vs. unstressed RMTg (Figure 7b), stressed RMTg vs stressed DRN (Figure 7c), and stressed RMTg vs stressed VTA (Figure 7d). Further, the Timberwolf, Turquoise Blue, and Silver modules reached a threshold of >1.5 in the Stressed DRN vs. Stressed RMTg and Stressed VTA vs. Stressed RMTg pathway comparisons (see Supplemental Code and Data). However, only the Gold and Silver modules demonstrated significant correlations between each gene’s centrality in the module and that gene’s Wald score. Interestingly, the Gold network genes tended to be decreased after stress in the RMTg pathway (i.e. downregulated) and were similarly downregulated in stressed RMTg compared to either the stressed DRN pathway or the stressed VTA pathway; we interpret this to mean that this gene module as a whole was downregulated in the stressed RMTg pathway.

**Figure 7.**
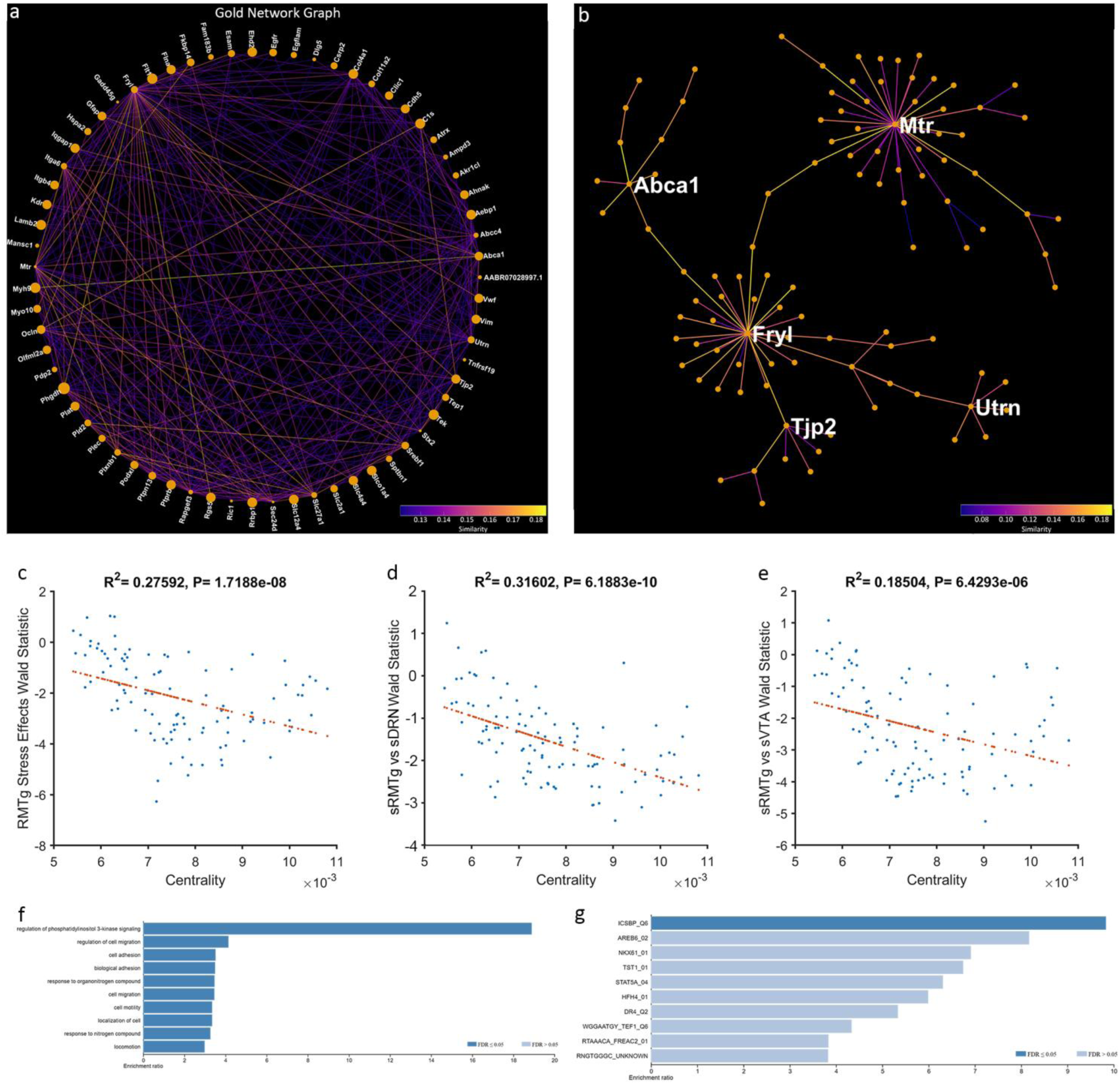
The Gold Network. **a)** Gold Gene Network projected onto a two-dimensional space using a circle layout. Line color represents pairwise gene expression similarity as calculated by WGCNA. Node size represents centrality to the network. **b**) Gold Network projected as a minimum spanning tree. The most central genes are labeled. **c**) Genes which are more downregulated with stress in the RMTg pathway are more central to the network (r2=0.276, p<0.001). **d**) Genes which are less highly expressed in the stressed RMTg than the stressed DRN pathway are more central to the network (r2=0.316, p<0.001). **e**) Genes which are less highly expressed in the stressed RMTg than the stressed VTA pathway are more central to the network (r2=0.185, p<0.001). **f**) Overrepresentation analysis of GO: Biological Process of the entire network. **g**) Overrepresentation analysis of transcription factor targets for the 25 most central genes to the network.

We next performed an overrepresentation analysis to determine if the genes in the Gold module were associated with identified gene sets using WebGestalt. While there was significant overrepresentation of genes from several pathways, the regulation of phosphoinositide 3 kinase (PI3K) signaling gene set was markedly overrepresented (enrichment ratio = 18.89; Figure 7e and Table 2). This gene set included Kdr, Pld2, Plxnb1, Ptpn13, and Tek. Furthermore, these genes are downregulated in the stressed RMTg pathway compared to the unstressed RMTg, the stressed VTA or the stressed DRN pathways. Table 2 shows the Wald statistic and q score for each gene in this set for each comparison. Additionally, when investigating the top 25 most central genes in this network, 6 of them are regulated by the transcription factor ICSBP (IRF8) (enrichment ratio = 9.8, FDR=0.01) (Figure 7f). This gene set included Col4a1, Fryl, Rrbp1, Slc12a4, Tek, and Tnfrsf19.

**Table 2.** PI3K Related Genes.

| Gene<br>Symbol | Gene<br>Name | Relative to sRMTg Pathway |  |  |  |  |  |
| --- | --- | --- | --- | --- | --- | --- | --- |
|  |  | usRMTg |  | sDRN |  | sVTA |  |
|  |  | Wald | Q | Wald | Q | Wald | Q |
| *Kdr* | kinase insert domain receptor | 4.829 | <0.001 | 2.133 | 0.301 | 3.871 | 0.008 |
| *Pld2* | phospholipase D2 | 2.096 | 0.543 | 2.577 | 0.234 | 2.569 | 0.121 |
| *Plxnb1* | plexin B1 | 2.512 | 0.331 | 2.139 | 0.301 | 2.426 | 0.147 |
| *Ptpn13* | protein tyrosine phosphatase,<br>non-receptor type 13 | 3.339 | 0.068 | 2.532 | 0.237 | 3.060 | 0.052 |
| *Tek* | TEK receptor tyrosine kinase | 4.067 | 0.008 | 2.128 | 0.303 | 4.389 | 0.002 |

## Discussion

In this study we investigated the impact of stress on RNAs actively undergoing translation in LHb neurons comprising the three major output pathways to VTA, DRN, and RMTg. We compared expression of individual genes between pathways and between unstressed and stressed conditions in a pairwise fashion. We found that GSEA and WGCNA revealed patterns in gene expression changes with stress that differed between these efferent LHb pathways. Whereas GSEA is based upon previously identified sets of functionally related genes, WGCNA identified a module of genes that was significantly differentially regulated in an unbiased fashion. In most cases, the gene sets or modules were downregulated in RMTg-projecting LHb neurons after stress but upregulated in DRN- and VTA-projecting neurons after stress.

When we compared the pathways to one another in a pairwise fashion in unstressed rats, there were relatively few differences in RNA expression; in particular, there were no dramatic phenotype-defining differences observed. A recent single cell RNAseq study in mice reached similar conclusions ^26^. While some LHb neurons express neuropeptides and perhaps GABA, nearly every neuron in the LHb is glutamatergic ^20,31,32^. Therefore, it is less surprising that the pathways are homogeneous at baseline and only change with stress. Our a priori hypothesis was that the neurons comprising these three pathways might be quite different because the pathways are highly segregated with few collaterals, but this was rejected; however, some differences between these pathways emerged following stress exposure.

Neurons projecting to different target regions are intermixed within subregions of the LHb yet have minimal collateralization between DRN, VTA, and RMTg. Given that the LHb neurons targeting these areas have similar phenotypes at baseline, perhaps the differences become apparent after stress because each pathway receives different input information during stress. The LHb receives inputs from myriad sources throughout the brain. Perhaps, stress-sensitive inputs preferentially synapse onto neurons projecting to specific targets, and differential activation leads to changes in their subsequent gene expression. Indeed, a recent paper found that CaMKII-expressing neurons projecting from the entopeduncular nucleus tended to synapse onto LHb neurons projecting to the VTA; whereas, those from the lateral hypothalamus and VTA tended to synapse onto neurons projecting to the DRN ^33^.

We found that RMTg-projecting neurons changed the most following stress. This was unexpected since we previously found that the pathway to the DRN was the most involved in adaptations to forced swim immobility ^4^. Chemogenetically inhibiting DRN-projecting neurons in between the first and second swim sessions decreased immobility and decreased perseverative seeking in the reward omission task. Conversely, activating these neurons increased perseverative seeking but did not further change immobility in the forced swim test. Inhibiting either of the other two pathways did not change behavior in any of the tests performed. Thus, we initially expected the pathway to the DRN to be the most affected by stress. One possible interpretation of these results is that inhibiting the LHb-RMTg pathway had no effect because the plasticity in this pathway already served to reduce its excitability, and inhibition with hM_4_Di had no additional impact. Supporting this interpretation, Proulx and colleagues found that acute optogenetic stimulation of RMTg-projecting LHb neurons transiently increased immobility during a single FST session ^34^, suggesting that it indeed is not maximally activated during forced swim alone. It is interesting to consider that, if the pathway from LHb to RMTg becomes less excitable due to changes in gene expression following forced swim stress, perhaps this leads to a relative increased activation of VTA and DRN by LHb.

The circuitry of the LHb and its major targets, RMTg, DRN, and VTA, is complex. LHb stimulation generally decreases dopaminergic activity within the VTA; although there is also evidence of direct excitation of dopamine neurons ^35^. Similarly, activation of the LHb has complex effects on the output of serotonin from raphe neurons; only high frequency stimulation increased serotonin release into striatum; this was further augmented by bicuculline injection into DRN ^6^. Indeed, lesions of the habenula ablate the increase in serotonin in the DRN during and after stress ^36^. LHb neurons project to both serotonergic and GABAergic neurons in DRN at roughly equal rates ^9^. While direct projections to VTA or DRN may either activate monoaminergic or GABAergic neurons, the projection to RMTg innervates only GABAergic neurons which in turn inhibit VTA and DRN ^37^. In this report we observed differential gene expression plasticity between neurons projecting to VTA and DRN on the one hand and RMTg on the other. Since the changes in RMTg-projecting neurons involved reduced expression of genes associated with excitability (such as in the PI3K signaling pathway), perhaps the overall balance of LHb stimulation of downstream GABAergic tone is reduced after repeated forced swim stress.

There is some diversity in the responses of LHb neurons to aversive stimuli. Both inescapable shock and reward omission increase firing in subpopulations of the LHb, exciting 30-50% of these neurons ^38,39^. Furthermore, roughly 10% of LHb neurons are actually inhibited by foot-shocks ^40^. Further, RMTg-projecting LHb neurons have shown increased excitability 24 hours to 14 days after cocaine exposure, but VTA-projecting neurons did not ^41,42^. Thus, it is possible that the observed stress-induced reduction in the excitability of LHb projections to RMTg disinhibits the VTA and DRN monoaminergic outputs. On the contrary, activation of LHb-RMTg neurons optogenetically interferes with reward motivated behavior and reduces activity in the FST ^34^. Perhaps the pathway to the RMTg becomes less active following an inescapable stressor via an allostatic adaptation in neuronal excitability.

Cerniauskas, et al. ^33^ characterized the effect of chronic mild stress on LHb neurons projecting to either the VTA/RMTg or the DRN. They observed greater excitability in neurons projecting to the VTA/RMTg in mice with the greatest composite depression-like behaviors. No such pattern was observed in the pathway to the DRN. Further, chemogenetic activation of LHb-VTA/RMTg neurons produced a depression-like phenotype in stress-naï ve mice. Conversely, DREADD-mediated inhibition of this pathway in stressed mice reduced their depression-like behaviors. Their data suggested that entopeduncular nucleus projections to LHb neurons that in turn project to VTA may be a critical functional circuit. Surgical targeting of VTA without hitting RMTg is difficult in mice, while these target areas can reliably be discriminated with retrograde strategies in rats ^4,7,43^, as we did in this study, so it is difficult to directly compare our results and theirs.

Our GSEA results suggest that gene expression in these pathways diverge after stress, with gene sets indicating RMTg-projecting LHb neurons changing in the opposite direction from those projecting to VTA or DRN. Of note, the gene sets related to oxidative phosphorylation and synaptic vesicle trafficking are upregulated with stress in the RMTg-projecting neurons and downregulated in the other two pathways; perhaps this is a compensatory adaptation to intense activation during stress. In several databases, gene sets related to extracellular matrix signaling were upregulated in the DRN and VTA pathways while these same gene sets were downregulated in the RMTg pathway following stress. The extracellular matrix signaling gene set was previously implicated in hippocampus following chronic stress ^44^, but it is difficult to predict what the impact of pathway-specific changes in extracellular matrix are in our results.

Unbiased topological analysis with WGCNA also indicated that the RMTg-projecting neurons diverged from the VTA- and DRN-projecting neurons. This was exemplified by the Gold module, which had an overrepresentation of downregulated genes associated with PI3K signaling after stress, found both via GO: Biological Process and the KEGG database. We propose that this reduction in PI3K-associated genes may reduce excitability of RMTg-projecting LHb neurons. PI3K signaling regulates anxiety-like behavior, may be involved in the antidepressant effects of deep brain stimulation, and is implicated in the mood-stabilizing effects of lithium in post-mortem analyses of patients with major depressive disorder ^45^. Upstream effectors of PI3K signaling are dysregulated in the LHb in a rat model of depression ^46^. Phosphorylation of PI3K is implicated in the antidepressant effects of baicalin in a mouse model of depression ^47^. We predict that downregulation of PI3K in the RMTg-projecting neurons after stress would be a compensatory adaptation that reduces their excitability, thereby disinhibiting the monoaminergic neurons in the DRN and VTA. Since we examined RNA actively undergoing translation in this study, the impact of these changes on protein production might take additional time to manifest. Future studies should examine the role of PI3K signaling in this pathway on resilience after stress exposure.

## Conclusions

In summary, RNA expression in the major efferent pathways from the LHb was more similar than anticipated at baseline, but stress led to divergent patterns of gene expression, particularly in RMTg-projecting neurons, and that might lead to altered balance of activity of these pathways during the response to stress.

## Methods

### Animals

For all experiments, male Sprague-Dawley rats (n=30, Charles River, Raleigh, NC) weighing 251-275 g were used. Rats were double-housed in a temperature- and humidity-controlled vivarium with a 14-10 light-dark cycle and allowed to acclimate to the facility for two weeks prior to surgery. All experiments were carried out during the light period. Food and water were freely available at all times. All experimental procedures were approved by the University of Washington Institutional Animal Care and Use Committee and were conducted in accordance to the guidelines of the ‘Principles of Laboratory Animal Care’ (NIH publication no. 86-23, 1996). 9 animals were excluded because viral-mediated gene expression of RiboTag was below the threshold.

### Experimental design

Our overarching experimental strategy was to use an intersectional viral vector approach to express RiboTag^28^ in LHb neurons selectively projecting to VTA, RMTg or DRN based using our recently published procedures ^4^. This was achieved through injection of Cre-dependent adeno-associated viral vectors (AAV8-hSyn-DIO-RiboTag) into LHb and a retrogradely transported canine adenovirus-2 expressing Cre (CAV2-Cre) into one of the three output regions. Separate groups of animals were used for each pathway to evaluate the effect of repeated swim stress on gene expression. All experimental procedures were approved by the University of Washington Institutional Animal Care and Use Committee and were conducted in accordance with National Institutes of Health (NIH) guidelines.

### Surgical Methods

For intersectional surgeries, anesthesia was induced with 5% isoflurane/95% oxygen and maintained at 1–3% isoflurane during the surgical procedure. Using a custom robotic stereotaxic instrument ^48^, 27 animals received AAV8-hSyn-DIO-RiboTag injected into LHb. Blunt 28g needles were inserted bilaterally at a 10° angle terminating at A/P −3.2, M/L ±0.7, and D/V −5.25 and 1 μl of AAV8-hSyn-DIO-RiboTag was injected at a rate of 0.2 μl/min. Nine animals received bilateral 1 μl injections of CAV2-Cre into the DRN at a 15° angle terminating at A/P −7.8, M/L ±0.23, and D/V −6.85. Nine animals received bilateral 1 μl injections of CAV2-Cre into the VTA at a 10° angle terminating at A/P −5.8, M/L ±0.6 and D/V −8.6. Nine animals received bilateral 1 μl injections of CAV2-Cre into the RMTg at a 10° angle terminating at A/P −7.6, M/L ±0.62 and D/V −8.5. After surgeries, rats were given meloxicam (0.2 mg/kg, s.c.) for pain management and monitored for at least 3 days. Accuracy of injection coordinates was confirmed by RTqPCR detection of RiboTag and Cre RNAs from LHb homogenate; these injection volumes and coordinates were optimized to produce selective transduction of LHb neurons with minimal expression adjacent regions. Rats recovered for three weeks post-surgery to give time for viral expression to occur. For “sham” control samples, rats were anesthetized and placed in the robotic stereotaxic instrument; however, no needle was inserted, nor any virus injected.

### Behavioral Methods

After three weeks of handling post-surgery, rats in the stressed grouping underwent a two-day forced swim protocol. Rats were placed in an inescapable 40cm tall x 20cm diameter Plexiglas cylinder filled with water (23±2C) to 30cm, a level deep enough to prevent them from standing on the bottom ^49^. Rats were stressed for 15 minutes on day one; 24 hours later, they were placed into the same chamber for five minutes. Rats in the unstressed group were handled as normal. Decapitation and tissue extraction occurred 3 hours following this protocol.

### RiboTag Extraction

RiboTag-associated RNA extraction as previously described ^29,50^.The LHb was extracted using a 4mm punch and homogenized in 2mL of supplemented homogenizing buffer [S-HB, 50 mM Tris-HCl, 100 mM KCl, 12 mM MgCl2, 1% NP40, 1 mM DTT, 1× Protease inhibitor cocktail (Sigma-Aldrich), 200 U/mL RNasin (Promega, Madison, WI), 100 µg/mL cyclohexamide (Sigma-Aldrich), 1 mg/mL heparin (APP Pharmaceuticals, Lake Zurich, IL)]. Samples were centrifuged at 4°C at 11,934 × g for 10 min, and supernatant was collected, reserving 50 µL (10%) as an input fraction. Mouse monoclonal HA-specific antibody (2.5 µL) (HA.11, ascites fluid; Covance, Princeton, NJ) was added to the remaining supernatant, and RiboTag-IP fractions were rotated at 4°C for 4 h. Protein A/G magnetic beads (200 µL) (Pierce) were washed with Homogenizing Buffer (HB 50 mM Tris-HCl, 100 mM KCl, 12 mM MgCl2, 1% NP40) prior to addition to the RiboTag-IP fraction and were rotated at 4°C overnight. The next day, RiboTag-IP fractions were placed on DynaMag-2 magnet (Life Technologies), and the bead pellet was washed 3 times for 15 min with high salt buffer (HSB; 50 mM Tris, 300 mM KCl, 12 mM MgCl2, 1% NP40, 1 mM DTT, and 100 µg/mL cyclohexamide) and placed on a rotator. After the final wash, HSB was removed and beads were re-suspended in 400 µL supplemented RLT buffer (10 µL β-mercaptoethanol/10 mL RLT Buffer) from the RNeasy Plus Micro Kit (Qiagen, Hilden, Germany) and vortexed vigorously. These samples were then placed back on the magnet and the RLT buffer was removed from the magnetic beads prior to RNA extraction. 350 µL supplemented RLT buffer was added to the Input Fraction prior to RNA extraction. RNA was extracted using Qiagen RNeasy Plus Micro kit according to package directions. RNA from both IP and input fractions were isolated using RNeasy Plus Micro Kit and eluted with 14-16µl of water. RNA concentration was measured using Quant-iT RiboGreen RNA Assay (ThermoFisher Cat. R11490, Waltham, MA). Total RNA yield for the RiboTag-IP fraction 36 ng (sham controls 12ng) and for the input fraction 196ng (sham controls 188ng).

### RNAseq Library Preparation

RNAseq libraries were prepared using SMARTer Stranded Total RNA-Seq Kit v2 – Pico Input Mammalian (Takara Bio USA, Inc. Cat. 635007, Mountain View, CA). 10ng of RNA or average equivalent volumes of “sham” control samples were used to generate the negative control samples. RNAseq libraries were submitted to Northwest Genomics Center at University of Washington (Seattle, WA) where library quality control was measured using a BioAnalyzer, library concentrations were measured using Qubit dsDNA HS Assay Kit (ThermoFisher), and then samples were normalized and pooled prior to cluster generation on HiSeq High Output for Paired-end reads. RNAseq libraries were sequenced on the HiSeq4000, Paired-end 75bp to sufficient read depth with PhiX spike-in controls (7%) (Illumina San Diego, CA).

### RT-qPCR Analysis

The remaining RNA was used to create cDNA libraries for qPCR using Superscript VILO Master Mix (ThermoFisher Cat. 11754050, Waltham, MA), and then cDNA libraries were diluted to a standard concentration before running the qPCR assay using Power Sybr Green on ViiA7 Real-Time PCR System (Thermo Fisher) or QuantStudio 5 Real-Time PCR System (Thermo Fisher). qPCR analysis was conducted using the standard curve method and normalized to four housekeeping genes (Gapdh, Ppia, Hprt, and Actinb). Normalized RSTQ data was analyzed using ANOVA with Bonferroni Post-Hoc.

### Bioinformatics

RNA sequencing was analyzed as previously described ^29,30^ with modification as noted below.

### Transcript Quantification and Quality Control

Raw fastq files were processed using multiple tools through the Galaxy platform26. Fastq files were inspected for quality using FastQC (Galaxy Version 0.7.0), and then passed to Salmon27 (Galaxy Version 0.8.2) for quantification of transcripts. The Salmon index was built using the protein coding transcriptome GRCm38-mm10.

### Non-Specific Immunoprecipitated RNA Subtraction

Due to the inherent issue of non-specific precipitation of immunoprecipitation (IP) procedures, we used a novel computational method for subtraction of non-specific RNA counts from the pathway specific RiboTag sample ^30^. Briefly, an IP was performed on an equivalent tissue sample from animals that did not receive the RiboTag virus (“sham” controls). RNA quantity from these samples was then quantified (mean ∼12ng) along with the RiboTag-expressing experimental IP samples (mean ∼36ng). Non-specific RNA contamination is considered to be a random sampling of the Input RNA, and the proportion of the IP sample that is contaminated is calculated from the true RNA quantity calculated for each sample. We capped the contamination level at 30% to prevent over-correction. All IP data presented in main manuscript have undergone this adjustment, but we also performed all bioinformatics on the raw IP files as well, which are available in the data archive. The adjusted bioinformatics are labeled “IP Minus Noise” while the raw files are labeled “IP”.

### Differential Expression Analysis

Differential gene expression was calculated using DESeq228 (Galaxy Version 2.11.39). All Salmon and DESeq2 settings were left default and our analysis pipeline is archived on our Galaxy server. Differential expression analysis was performed on the IP and Input samples. To determine pathway specific gene enrichment, all IP samples were compared to pooled Input samples. For all other test of differential expression, IP samples were compared to IP samples. To determine the pathway expression at baseline, the unstressed DRN pathway was compared to the unstressed RMTg or VTA pathways and the unstressed RMTg pathway was compared to the unstressed VTA pathway. To determine the effects of stress between each pathway, the stressed DRN pathway was compared to the stressed RMTg or VTA pathways and the stressed RMTg pathway was compared to the stressed VTA pathway. To determine the effects of stress within each pathway, the stressed DRN pathway was compared to the unstressed DRN pathway, stressed RMTg to unstressed RMTg, and stressed VTA to unstressed VTA. For all comparisons, a positive Wald statistic means that a gene is expressed more in the first group as compared to the second group. An FDR of q = 0.10 was used as the threshold for differential expression throughout the manuscript.

### Gene Set Enrichment Analysis (GSEA)

Wald statics generated by DESeq2 were used as the ranking variable for gene set enrichment analysis (GSEA). The Wald statistic has a benefit over log2(fold change) as it incorporates an estimate of variance, and ultimately is used to determine significance in differential expression analysis. All genes with reliable statistical comparisons (those not filtered by DeSeq2) were entered into WebGestalt 201929 ^51^ and we performed GSEA on all pertinent comparisons (Stressed vs Unstressed for each pathway, pairwise comparisons between the three stressed pathways, and pairwise comparisons between the three unstressed pathways). Gene sets used include three Gene Ontology (GO) classes: Biological Process, Molecular Function, and Cellular Component, as well as Kyoto Encyclopedia of Genes and Genomes (KEGG) pathways, Panther, Transcription Factor Targets, and MicroRNA Targets. All advanced parameters were left default except for significance level, which was set to FDR = 0.1.

### Weighted Gene Co-expression Network Analysis (WGCNA)

Topological overlap matrix for IP samples were generated, and module clustering was accomplished using the WGCNA ^52^ package for R ^53^. Briefly, the TPM matrix for each group was filtered to remove zero-variance genes, and a signed adjacency matrix was generated using “bicor” as the correlation function. From this a signed topological overlap matrix was generated, followed by a dissimilarity topological overlap matrix. Finally, module membership was assigned using a dynamic tree cut. The complete code used to run WGCNA are available in the data archive. Post processing of the topological overlap matrix was completed with a custom object based Matlab class structure that we released previously ^30^. The WGCNA class structure can be accessed on https://github.com/levinsmr/WGCNA and the code used to generate all of the WGCNA figures in this manuscript is available in the Supplementary Code for use as an example. When the average Wald score of a module was >2.0, we performed a secondary overrepresentation analysis of the component genes to evaluate for overrepresentation of genes associated with particular biological functions with WebGestalt.

## Code and Data Availability

Code and data used to generate this manuscript are available for direct download. RNAseq files and the RNAseq pipeline are available on our Galaxy Server. Code used for figure generation are available in the **Supplementary Code**.

## Acknowledgments

supported by MH106532 (JFN), NS099578 (MRL), and DA007278 (KRC).

## Author Contributions

MRL – Conceptualization, methodology, formal analysis, investigation, writing – original draft, visualization

KRC – methodology, software, visualization, writing – review & editing

RM – methodology, software

AJL – conceptualization, methodology

JFN – Conceptualization, resources, supervision, funding acquisition, project administration, writing – review & editing

---

**Supplemental Figure 1.**
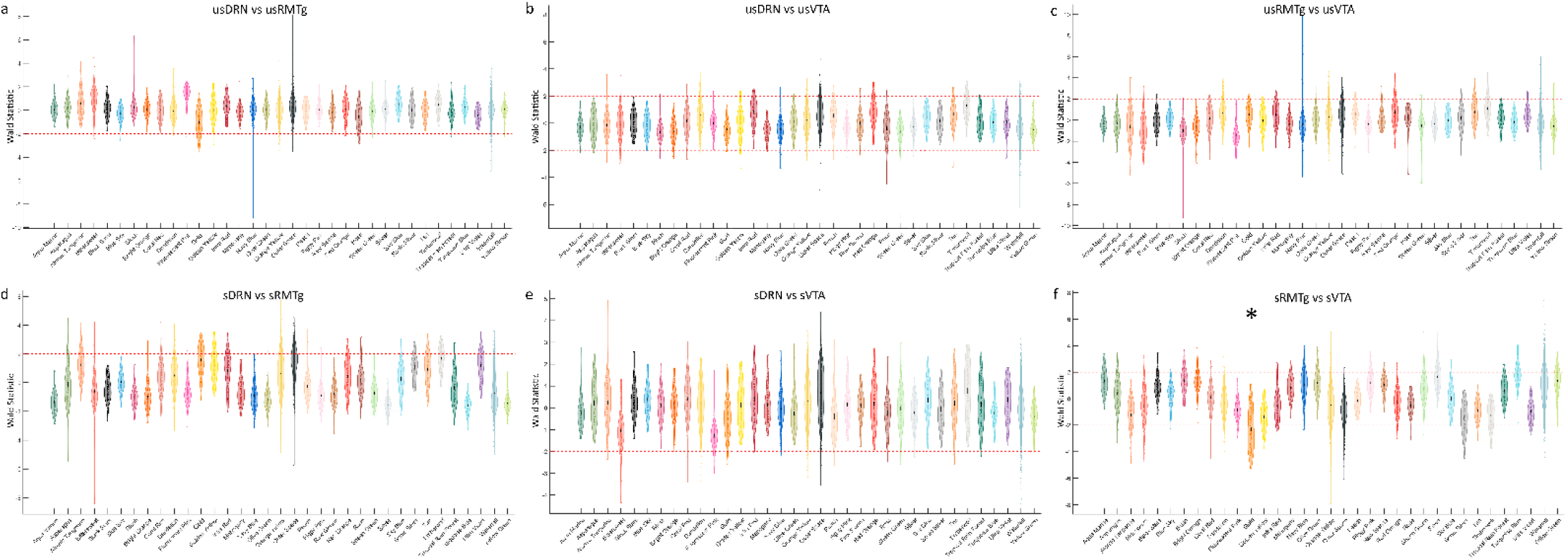
The Wald score for each gene within the 35 modules assigned by WGCNA for remaining pairwise comparisons. **a)** Unstressed DRN pathway compared to Unstressed RMTg pathway. **b)** Unstressed DRN pathway compared to Unstressed VTA pathway. **c)** Unstressed RMTg pathway compared to Unstressed VTA pathway. **d**) Stressed DRN pathway compared to Stressed RMTg pathway. **e)** Stressed DRN pathway compared to Stressed VTA pathway. **f)** Stressed RMTg pathway compared to Stressed VTA pathway.

**Supplemental Figure 2.**
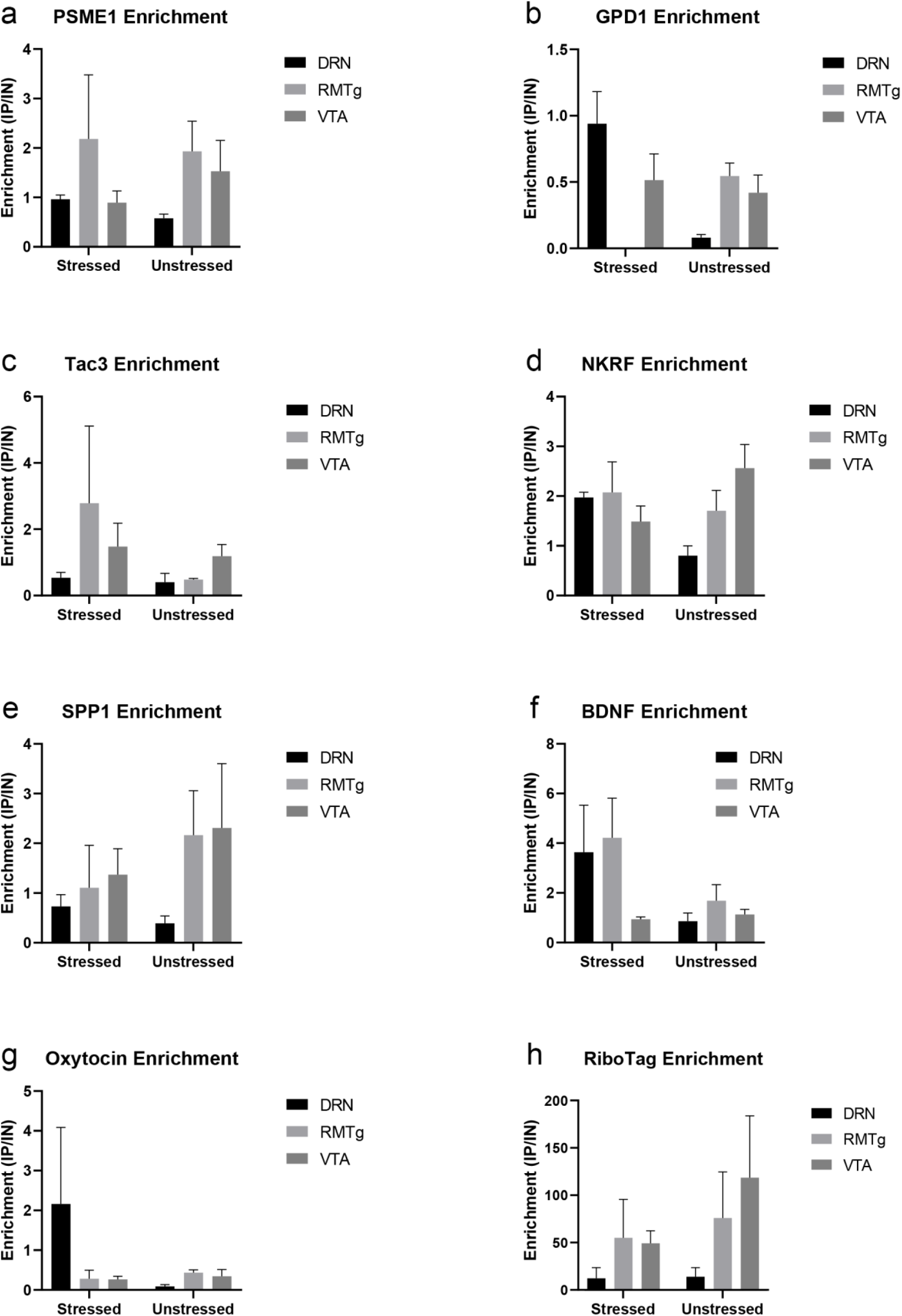
RTqPCR validation of gene targets. **a**) Enrichment of PSME1 in each pathway. **b**) Enrichment of GPD1 in each pathway. **c**) Enrichment of Tac3 in each pathway. d) Enrichment of NKRF in each pathway. **e)** Enrichment of SPP1 in each pathway. **f)** Enrichment of BDNF in each pathway. **g)** Enrichment of Oxytocin in each pathway. h) Enrichment of RiboTag in each pathway.

**Supplemental Figure 3.**
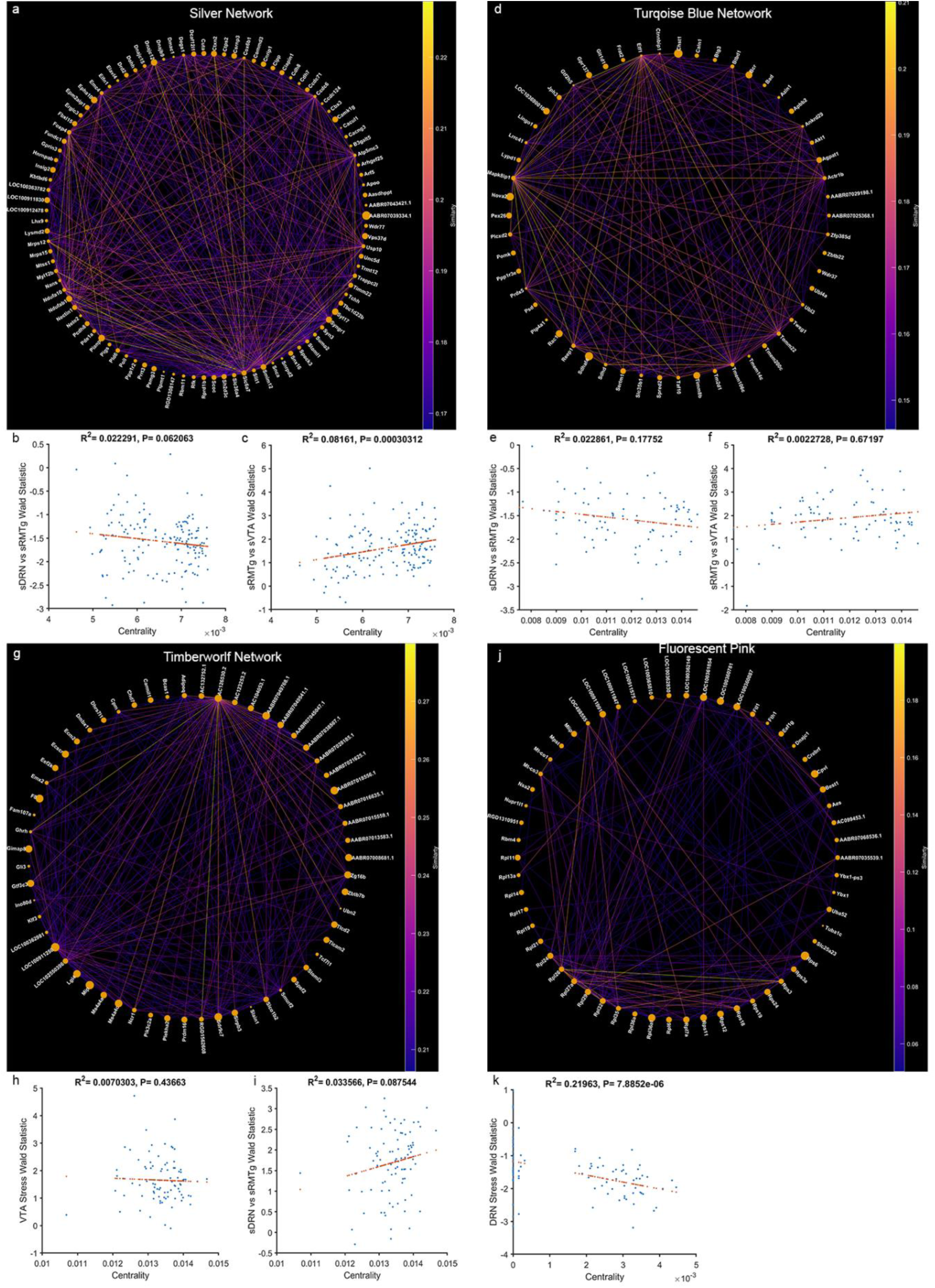
Networks passing the ±1.5 threshold. **a**) Silver network circular graph. **b**) Correlation of centrality of gene with the sDRN vs sRMTg Wald statistic for the Silver network. **c**) Correlation of centrality of gene with the sRMTg vs sVTA Wald statistic for the Silver network. **d**) Turquoise Blue network circular graph. **e**) Correlation of centrality of gene with the sDRN vs sRMTg Wald statistic for the Turquoise Blue network. **f)** Correlation of centrality of gene with the sRMTg vs sVTA Wald statistic for the Silver network. **g**) Timberwolf network circular graph. **h**) Correlation of centrality of gene with the VTA Stress Wald statistic for the Timberwolf network. **i**) Correlation of centrality of gene with the sDRN vs sRMTg Wald statistic for the Timberwolf network. j) Fluorescent Pink network circular graph. **k)** Correlation of centrality of gene with the DRN Stress Wald statistic for the Fluorescent Pink network.

## References

1 Williams, L. M. Precision psychiatry: a neural circuit taxonomy for depression and anxiety. Lancet Psychiatry 3, 472–480, doi: 10.1016/S2215-0366(15)00579-9 (2016).

2 Geisler, S. & Trimble, M. The lateral habenula: no longer neglected. CNS spectrums 13, 484–489, doi: 10.1017/s1092852900016710 (2008).

3 Brinschwitz, K. et al. Glutamatergic axons from the lateral habenula mainly terminate on GABAergic neurons of the ventral midbrain. Neuroscience 168, 463–476, doi: 10.1016/j.neuroscience.2010.03.050 (2010).

4 Coffey, K. R., Marx, R. G., Vo, E. K., Nair, S. G. & Neumaier, J. F. Chemogenetic inhibition of lateral habenula projections to the dorsal raphe nucleus reduces passive coping and perseverative reward seeking in rats. Neuropsychopharmacology, doi: 10.1038/s41386-020-0616-0 (2020).

5 Quina, L. A. et al. Efferent pathways of the mouse lateral habenula. J Comp Neurol 523, 32–60, doi: 10.1002/cne.23662 (2015).

6 Kalen, P., Strecker, R. E., Rosengren, E. & Bjorklund, A. Regulation of striatal serotonin release by the lateral habenula-dorsal raphe pathway in the rat as demonstrated by in vivo microdialysis: role of excitatory amino acids and GABA. Brain research 492, 187–202, doi: 10.1016/0006-8993(89)90901-3 (1989).

7 Goncalves, L., Sego, C. & Metzger, M. Differential projections from the lateral habenula to the rostromedial tegmental nucleus and ventral tegmental area in the rat. J Comp Neurol 520, 1278–1300, doi: 10.1002/cne.22787 (2012).

8 Bernard, R. & Veh, R. W. Individual neurons in the rat lateral habenular complex project mostly to the dopaminergic ventral tegmental area or to the serotonergic raphe nuclei. J Comp Neurol 520, 2545–2558, doi: 10.1002/cne.23080 (2012).

9 Weissbourd, B. et al. Presynaptic partners of dorsal raphe serotonergic and GABAergic neurons. Neuron 83, 645–662, doi: 10.1016/j.neuron.2014.06.024 (2014).

10 Ogawa, S. K., Cohen, J. Y., Hwang, D., Uchida, N. & Watabe-Uchida, M. Organization of monosynaptic inputs to the serotonin and dopamine neuromodulatory systems. Cell Rep 8, 1105–1118, doi: 10.1016/j.celrep.2014.06.042 (2014).

11 Kowski, A. B., Veh, R. W. & Weiss, T. Dopaminergic activation excites rat lateral habenular neurons in vivo. Neuroscience 161, 1154–1165, doi: 10.1016/j.neuroscience.2009.04.026 (2009).

12 Xie, G. et al. Serotonin modulates glutamatergic transmission to neurons in the lateral habenula. Sci Rep 6, 23798, doi: 10.1038/srep23798 (2016).

13 Zhang, H. et al. Dorsal raphe projection inhibits the excitatory inputs on lateral habenula and alleviates depressive behaviors in rats. Brain Struct Funct 223, 2243–2258, doi: 10.1007/s00429-018-1623-3 (2018).

14 Shen, X., Ruan, X. & Zhao, H. Stimulation of midbrain dopaminergic structures modifies firing rates of rat lateral habenula neurons. PloS one 7, e34323, doi: 10.1371/journal.pone.0034323 (2012).

15 Han, L. N. et al. Activation of serotonin(2C) receptors in the lateral habenular nucleus increases the expression of depression-related behaviors in the hemiparkinsonian rat. Neuropharmacology 93, 68–79, doi: 10.1016/j.neuropharm.2015.01.024 (2015).

16 Lee, E. H. & Huang, S. L. Role of lateral habenula in the regulation of exploratory behavior and its relationship to stress in rats. Behavioural brain research 30, 265–271, doi: 10.1016/0166-4328(88)90169-6 (1988).

17 Browne, C. A., Hammack, R. & Lucki, I. Dysregulation of the Lateral Habenula in Major Depressive Disorder. Front Synaptic Neurosci 10, 46, doi: 10.3389/fnsyn.2018.00046 (2018).

18 Wirtshafter, D., Asin, K. E. & Pitzer, M. R. Dopamine agonists and stress produce different patterns of Fos-like immunoreactivity in the lateral habenula. Brain research 633, 21–26, doi: 10.1016/0006-8993(94)91517-2 (1994).

19 Ootsuka, Y. & Mohammed, M. Activation of the habenula complex evokes autonomic physiological responses similar to those associated with emotional stress. Physiol Rep 3, doi: 10.14814/phy2.12297 (2015).

20 Stamatakis, A. M. & Stuber, G. D. Activation of lateral habenula inputs to the ventral midbrain promotes behavioral avoidance. Nature neuroscience 15, 1105–1107, doi: 10.1038/nn.3145 (2012).

21 Gill, M. J., Ghee, S. M., Harper, S. M. & See, R. E. Inactivation of the lateral habenula reduces anxiogenic behavior and cocaine seeking under conditions of heightened stress. Pharmacology, biochemistry, and behavior 111, 24–29, doi: 10.1016/j.pbb.2013.08.002 (2013).

22 Zhang, Q. et al. Lateral habenula as a link between thyroid and serotoninergic system modiates depressive symptoms in hypothyroidism rats. Brain research bulletin 124, 198–205, doi: 10.1016/j.brainresbull.2016.05.007 (2016).

23 Luo, X. F. et al. Lateral habenula as a link between dopaminergic and serotonergic systems contributes to depressive symptoms in Parkinson’s disease. Brain Res. Bull. 110, 40–46, doi: 10.1016/j.brainresbull.2014.11.006 (2015).

24 Nair, S. G., Strand, N. S. & Neumaier, J. F. DREADDing the lateral habenula: a review of methodological approaches for studying lateral habenula function. Brain Res. 1511, 93–101, doi: 10.1016/j.brainres.2012.10.011 (2013).

25 Wagner, F., French, L. & Veh, R. W. Transcriptomic-anatomic analysis of the mouse habenula uncovers a high molecular heterogeneity among neurons in the lateral complex, while gene expression in the medial complex largely obeys subnuclear boundaries. Brain Struct Funct 221, 39–58, doi: 10.1007/s00429-014-0891-9 (2016).

26 Wallace, M. L. et al. Anatomical and single-cell transcriptional profiling of the murine habenular complex. Elife 9, doi: 10.7554/eLife.51271 (2020).

27 Hashikawa, Y. et al. Transcriptional and Spatial Resolution of Cell Types in the Mammalian Habenula. Neuron 106, 1–16, doi: 10.1016/j.neuron.2020.03.011 (2020).

28 Sanz, E. et al. Cell-type-specific isolation of ribosome-associated mRNA from complex tissues. Proc. Natl. Acad. Sci. U. S. A. 106, 13939–13944, doi: 10.1073/pnas.0907143106 (2009).

29 Lesiak, A. J. et al. Sequencing the Serotonergic Neuron Translatome Reveals a New Role for Fkbp5 in Stress. Molecular psychiatry (2020).

30 Coffey, K. R. et al. RiboTag-Seq Reveals a Compensatory cAMP Responsive Gene Network in Striatal Microglia Induced by Morphine Withdrawal. bioRxiv, doi: 10.1101/2020.02.10.942953 (2020).

31 Zuo, W. et al. Ethanol potentiates both GABAergic and glutamatergic signaling in the lateral habenula. Neuropharmacology 113, 178–187, doi: 10.1016/j.neuropharm.2016.09.026 (2017).

32 Aizawa, H., Kobayashi, M., Tanaka, S., Fukai, T. & Okamoto, H. Molecular characterization of the subnuclei in rat habenula. J. Comp. Neurol. 520, 4051–4066, doi: 10.1002/cne.23167 (2012).

33 Cerniauskas, I. et al. Chronic Stress Induces Activity, Synaptic, and Transcriptional Remodeling of the Lateral Habenula Associated with Deficits in Motivated Behaviors. Neuron 104, 899–915 e898, doi: 10.1016/j.neuron.2019.09.005 (2019).

34 Proulx, C. D. et al. A neural pathway controlling motivation to exert effort. Proc. Natl. Acad. Sci. U. S. A. 115, 5792–5797, doi: 10.1073/pnas.1801837115 (2018).

35 Brown, P. L. & Shepard, P. D. Functional evidence for a direct excitatory projection from the lateral habenula to the ventral tegmental area in the rat. Journal of neurophysiology 116, 1161–1174, doi: 10.1152/jn.00305.2016 (2016).

36 Amat, J. et al. The role of the habenular complex in the elevation of dorsal raphe nucleus serotonin and the changes in the behavioral responses produced by uncontrollable stress. Brain research 917, 118–126, doi: 10.1016/s0006-8993(01)02934-1 (2001).

37 Yang, Y., Wang, H., Hu, J. & Hu, H. Lateral habenula in the pathophysiology of depression. Current opinion in neurobiology 48, 90–96, doi: 10.1016/j.conb.2017.10.024 (2018).

38 Lecca, S. et al. Aversive stimuli drive hypothalamus-to-habenula excitation to promote escape behavior. Elife 6, doi: 10.7554/eLife.30697 (2017).

39 Shabel, S. J., Wang, C., Monk, B., Aronson, S. & Malinow, R. Stress transforms lateral habenula reward responses into punishment signals. Proc. Natl. Acad. Sci. U. S. A. 116, 12488–12493, doi: 10.1073/pnas.1903334116 (2019).

40 Congiu, M., Trusel, M., Pistis, M., Mameli, M. & Lecca, S. Opposite responses to aversive stimuli in lateral habenula neurons. The European journal of neuroscience 50, 2921–2930, doi: 10.1111/ejn.14400 (2019).

41 Meye, F. J. et al. Cocaine-evoked negative symptoms require AMPA receptor trafficking in the lateral habenula. Nature neuroscience 18, 376–378, doi: 10.1038/nn.3923 (2015).

42 Maroteaux, M. & Mameli, M. Cocaine evokes projection-specific synaptic plasticity of lateral habenula neurons. The Journal of neuroscience: the official journal of the Society for Neuroscience 32, 12641–12646, doi: 10.1523/JNEUROSCI.2405-12.2012 (2012).

43 Fu, R. et al. Low-dose ethanol excites lateral habenula neurons projecting to VTA, RMTg, and raphe. Int J Physiol Pathophysiol Pharmacol 9, 217–230 (2017).

44 Li, X. H. et al. Gene expression profile of the hippocampus of rats subjected to chronic immobilization stress. PloS one 8, e57621, doi: 10.1371/journal.pone.0057621 (2013).

45 Voleti, B. & Duman, R. S. The roles of neurotrophic factor and Wnt signaling in depression. Clin Pharmacol Ther 91, 333–338, doi: 10.1038/clpt.2011.296 (2012).

46 Christensen, T., Jensen, L., Bouzinova, E. V. & Wiborg, O. Molecular profiling of the lateral habenula in a rat model of depression. PloS one 8, e80666, doi: 10.1371/journal.pone.0080666 (2013).

47 Guo, L. T. et al. Baicalin ameliorates neuroinflammation-induced depressive-like behavior through inhibition of toll-like receptor 4 expression via the PI3K/AKT/FoxO1 pathway. J Neuroinflammation 16, 95, doi: 10.1186/s12974-019-1474-8 (2019).

48 Coffey, K. R., Barker, D. J., Ma, S. & West, M. O. Building an open-source robotic stereotaxic instrument. J Vis Exp, e51006, doi: 10.3791/51006 (2013).

49 Clark, M. S. et al. Overexpression of 5-HT1B receptor in dorsal raphe nucleus using Herpes Simplex Virus gene transfer increases anxiety behavior after inescapable stress. The Journal of neuroscience: the official journal of the Society for Neuroscience 22, 4550–4562, doi: 10.1523/JNEUROSCI.22-11-04550.2002 (2002).

50 Lesiak, A. J. & Neumaier, J. F. RiboTag: Not Lost in Translation. Neuropsychopharmacology: official publication of the American College of Neuropsychopharmacology 41, 374–376, doi: 10.1038/npp.2015.262 (2016).

51 Liao, Y., Wang, J., Jaehnig, E. J., Shi, Z. & Zhang, B. WebGestalt 2019: gene set analysis toolkit with revamped UIs and APIs. Nucleic Acids Res. 47, W199–W205, doi: 10.1093/nar/gkz401 (2019).

52 Langfelder, P. & Horvath, S. WGCNA: an R package for weighted correlation network analysis. BMC Bioinformatics 9, 559, doi: 10.1186/1471-2105-9-559 (2008).

53 R: A language and environment for statistical computing (R Foundation for Statistical Computing, Vienna, Austria, 2020).

